# The LINC complex inhibits excessive chromatin repression

**DOI:** 10.1101/2022.02.09.479725

**Authors:** Daria Amiad Pavlov, CP Unnikannan, Dana Lorber, Gaurav Bajpai, Tsviya Olender, Elizabeth Stoops, Adriana Reuveny, Samuel Safran, Talila Volk

**Affiliations:** Department of Molecular Genetics, Weizmann Institute of Science, Rehovot, Israel, DAP current address: Department of Physiology, University of Pennsylvania Perelman School of Medicine; Bayer AG, Leverkusen 51368, Germany; Department of Chemical and Biological Physics, Weizmann Institute of Science, Rehovot

## Abstract

We show here that the Linker of Nucleoskeleton and Cytoskeleton (LINC) complex is needed to minimize chromatin repression. The genomic binding profile of Polycomb, Heterochromatin Protein1 (HP1a) repressors, and of RNA-Pol II were studied in *Drosophila* larval muscles lacking functional LINC complex. A significant increase in the binding of Polycomb, and parallel reduction of RNA-Pol-II binding to a set of muscle genes was observed. Consistently, enhanced tri-methylated H3K9 and H3K27 repressive modifications, and reduced chromatin activation by H3K9 acetylation were found. Furthermore, larger tri-methylated H3K27me3 repressive clusters, and chromatin redistribution from the nuclear periphery towards nuclear center, were detected in live LINC mutant larval muscles. Computer simulation indicated that the observed dissociation of the chromatin from the nuclear envelope promotes growth of tri-methylated H3K27 repressive clusters. Thus, we suggest that by promoting chromatin-nuclear envelope binding, the LINC complex restricts the size of repressive H3K27 tri-methylated clusters, thereby limiting the binding of Polycomb transcription repressor, directing robust transcription in muscle fibers.

## Introduction

The Linker of Nucleoskeleton and Cytoskeleton (LINC) complex, implicated in mechanical coupling between the cytoplasm and nucleoplasm, has been proposed to control chromatin organization and the epigenetic state of chromatin (Kirby and Lammerding, 2018; Lityagina and Dobreva, 2021; Poulet et al., 2017; Wagh et al., 2021). This complex consists of Nesprin proteins whose membrane-KASH domain is inserted into the outer nuclear membrane and their N-terminus associates with various cytoskeletal components. At the perinuclear space Nesprin tetramers bind covalently to SUN domain tetramers which further associate with the nuclear lamina and with chromatin (Chang et al., 2015; Razafsky and Hodzic, 2009; Sosa et al., 2012; Tapley and Starr, 2013; Wallrath et al., 2016). The functional contribution of the LINC complex for human health is displayed by numerous diseases associated with mutations in genes coding for components of the LINC complex, including Emery Dreifuss Muscular Dystrophy (EDMD), Arthrogryposis, Cerebral ataxia, Dilated Cardiomyopathy (DCM) and others (Cartwright and Karakesisoglou, 2014; Méjat and Misteli, 2010; Puckelwartz et al., 2009; Horn, 2014). Whereas the primary cause of LINC-associated diseases is not entirely clear, lack of functional LINC complex has been recently suggested to alter chromatin organization resulting in defects with gene transcription, and aberrant tissue function. For example, depletion of Nesprin-3 leads to nucleus collapse and loss of genome organization in adult rat cardiomyocytes (Heffler et al., 2019), LINC disruption in embryonic cardiomyocytes led to global chromatin and H3K9me3 rearrangement (Seelbinder et al., 2021), plants deficient for components of the LINC complex, such as KASH (wifi) and SUN (sun1 sun4 sun5 triple mutant) show altered nuclear shape, increased distance of chromocenters from the nuclear periphery, altered heterochromatin organization and reactivation of transcriptionally silent repetitive sequences (Poulet et al., 2017). Likewise, mouse keratinocytes lacking SUN proteins exhibited precocious epidermal differentiation and loss of repressive chromatin H3K27me3 mark on differentiation-specific genes (Carley et al., 2021). In oligodendrocytes silencing of the Nesprin Syne1 gene resulted in aberrant histone marks, chromatin reorganization and impaired gene transcription (Hernandez et al., 2016). Furthermore, in *S. cerevisiae* Mps3, a SUN homologue is involved in the recruitment of heterochromatic sequences such as telomeric repeats to the nuclear envelope (NE), an essential process needed for spindle formation in the course of chromosome segregation (Ghosh et al., 2012). These studies implied that the LINC complex function associates with regulation of the epigenetic state of the chromatin, however the underlying mechanism is currently unclear. Moreover, most of these studies were performed either on cells in culture conditions, or on cells that were not fully differentiated. The contribution of the LINC complex to fully differentiated, non-dividing cells, where chromatin landscape has been stabilized, is yet to be elucidated.

Here, we show that the LINC complex is needed to minimize chromatin repression. Previously we showed that the LINC complex is required for proper larval muscle contractions, and for nuclear position along the entire muscle fiber (Elhanany-Tamir et al., 2012; Wang et al., 2015), as well as for synchronization of myonuclear mechanical dynamics (Lorber et al., 2020). In the present study we analyzed the genomic binding profile of the transcription repressors Polycomb and HP1a, as well as that of RNA-Pol II, and quantified the levels of repressive epigenetic chromatin marks, tri-methylated H3K27 and H3K9, and active H3K9ac chromatin mark, in *Drosophila* larval muscles lacking functional LINC complex. We found that differentiated muscles lacking functional LINC complex exhibited increased epigenetic repression that correlated with enhanced genome binding of Polycomb and HP1a, together with reduced binding of RNA-Pol II to a set of essential muscle genes. The increased chromatin repression observed in the *SUN/koi* mutant muscle nuclei correlated with increased size of tri-methylated H3K27 chromatin clusters, and with partial translocation of chromatin from the periphery to nuclear center quantified from live 3D imaging. Simulations of the 3D distribution of tri-methylated H3K27 sites indicated that reduced binding of chromatin to the nuclear lamina leads to increased repressive cluster size. These results suggest that the enhanced chromatin repression observed in the LINC mutants, was induced by weakening of chromatin binding to the nuclear lamina. Altogether our results imply that by coupling between the cytoskeleton and the nucleoskeleton the LINC complex restrains gene repression in mature muscle fibers.

## Results

### Genome-wide Polycomb, HP1a and RNA-Pol II binding profiles reveal increased repression and reduced activation in *SUN/koi* mutant muscle fibers

Although the LINC complex has been implicated in the regulation of epigenetic state, chromatin organization and overall gene-expression (Wagh et al., 2021), it is not clear whether and how LINC mediated mechanotransduction is involved in the downstream genome-wide chromatin repression and activation. Here, we investigated the contribution of the LINC complex to the epigenetic chromatin state by analyzing the phenotype of *SUN/koi* mutant muscles in *Drosophila* 3^rd^ instar larvae. Whereas the two Nesprin-like genes of *Drosophila* have additional functions in cells, due to non-KASH containing isoforms, *SUN/koi*, the only SUN representative in *Drosophila* does not appear to exhibit LINC-unrelated functions (Elhanany-Tamir et al., 2012; Titlow et al., 2020). We therefore investigated changes to genome-wide chromatin repression and activation states in the fully differentiated muscles of *SUN/koi* mutants in-vivo, using the Targeted DamID (TaDa) technique that allows tissue-specific profiling of DNA-binding proteins, in the intact living organism (Southall et al., 2013). Here we compared the DNA-binding profiles in *SUN/koi* mutated and control muscle fibers of two chromatin factors: (1) Polycomb, the H3K27me3 reader representative of the polycomb-group associated heterochromatin, (2) HP1a, the H3K9me3 reader representative of the HP1a-associated heterochromatin, and in addition, the profile of RNA-Pol II subunit representative of actively transcribed euchromatin. Polycomb, HP1a, and RNA-pol II fused to the DNA adenine methyltransferase (Dam) were driven to *Drosophila* larval muscles using Mef2-GAL4 driver, in a temporally controlled manner (by combined expression of the temperature sensitive inhibitor Gal80^ts^), inducing expression of the Dam-fusion proteins only at the 3^rd^ instar larval stage, when muscles are mature and fully differentiated. The experiment included N=3 independent replicates of Dam-Polycomb, Dam-HP1a, Dam-RNA Pol II, and Dam only (as reference), for control and for *SUN*/*koi* nutant genotypes, with n=25 larvae in each group. Following 10 hours of Dam expression, larvae were dissected, DNA was extracted and DpnI digested at the Dam methylated GATC sites. Further amplification, processing, and bioinformatics analysis were performed as described (Marshall and Brand, 2015).

To compare gene binding profiles between *SUN*/*koi* and control muscle fibers, we performed a PCA regression analysis on Dam-Polycomb, Dam-HP1a, and Dam-Pol II, normalized to Dam binding alone (Figure 1A). The cut-off criteria for significant change in gene binding between *SUN/koi* and control was set to false discovery rate (FDR) < 0.05, z-score > 1.96, and GATC sites > 1. Gray dots in Figure 1A represent genes with significantly altered binding, such that genes below the regression line represent increased binding in the *SUN/koi* mutant, and genes above the regression line represent decreased binding in *SUN/koi*, compared to control. Overall, we detected significantly altered binding to 148 genes in the Polycomb group, 173 genes in the HP1a group and 206 genes in the RNA-Pol II group. Note the robust increase in Polycomb binding observed in the *SUN/koi* mutant, where 143 out of 148 significant genes showed enhanced Polycomb occupancy. Notably, there was only 0.5 % and 2.4% overlap between the Polycomb-RNA-Pol II, and HP1a-RNA-Pol II groups, respectively, and 6.3% overlap between the Polycomb-HP1a groups. Figure 1B illustrates representative examples of non-overlapping occupancy of Dam-Polycomb, Dam-HP1a, and Dam-Pol II in 3 genes expressed in the larval muscle fibers, namely alphaTub84B (left), Nup62 (middle), and Act57B (right) genes. Interestingly, each of these genes showed altered occupancy in *SUN/koi* mutant muscles for only one chromatin factor (red boxes) with increased Polycomb binding to alphaTub84B, increased HP1a binding to Nup62, and decreased RNA-Pol II binding to the Act57B gene. The corresponding Zpca scores are listed for each gene on Figure 1B.

**Figure 1.**
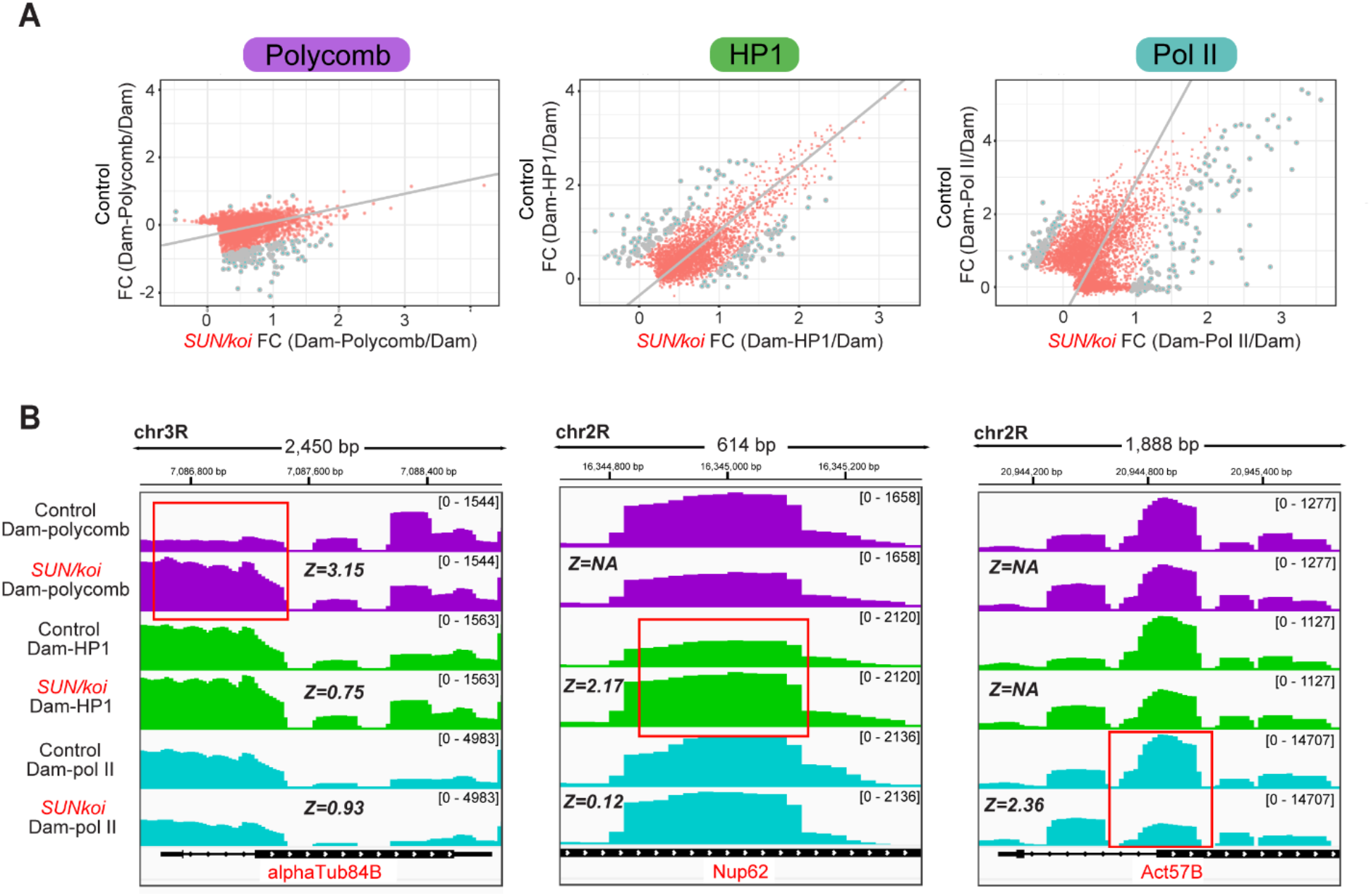
Increased Polycomb and altered HP1a and RNA-Pol II occupancies in LINC mutated muscle fibers. **(A)** PCA regression on Dam-Polycomb (left), Dam-HP1a (middle), Dam-Pol II (right) gene occupancy, normalized to Dam alone, in control versus *SUN*/*koi* mutated muscle fibers. Gray dots represent genes with significantly altered binding profile when cut-off criteria are set to FDR<0.05, Z-score >1.96, GATC sites>1. **(B)** Dam-Polycomb, Dam-HP1a, and Dam-Pol II, occupancy, normalized to Dam alone, for the alphaTub84B (left), Nup62 (middle), and Act57B (right) genes. Each gene presents altered occupancy in only one chromatin factor upon *SUN*/*koi* mutation (highlighted with red box) and the corresponding Zpca scores are listed for each gene.

We further paralleled genome wide changes in Polycomb, HP1a and RNA-Pol II occupancy in the *SUN/koi* mutant muscles by comparing the significantly altered genes in each group. Figure 2A depicts a heatmap of (*SUN/koi* versus control) fold change occupancy, for each chromatin factor, such that red indicates increased occupancy and blue indicates decreased occupancy as observed in *SUN/koi* mutant muscles. Non-significant genes are labeled in gray. Gene clustering was defined by k-means, and the number of clusters was estimated with Gap statistics. Overall, we observe minimal overlap between the group of genes that showed altered binding to the three factors, namely, Polycomb, HP1a, and RNA-Pol II, as the majority of genes colored in each group are non-significant (gray) in the others. Cluster 1 in Figure 2A includes genes predicted to undergo decreased transcription. It contains the genes that showed increased Polycomb occupancy, increased HP1a binding, and decreased RNA-Pol II binding. Cluster 2 also includes a group of genes predicted to undergo reduced transcription due to decreased RNA-Pol II binding. These genes partially overlapped with reduced HP1a occupancy genes, possibly representing genes that normally are positively regulated by HP1a (Kwon and Workman, 2011). Cluster 3 includes 3 non-overlapping groups of genes: those exhibited increased RNA-Pol II binding, and therefore expected to be upregulated, and two groups of genes, which showed increased, or decreased occupancy by HP1a. The latter two groups might represent genes that are normally either repressed, or activated by HP1a, and in the mutant are affected in opposing direction. Further, the three clusters showed non-overlapping functionalities according to GO analysis (Figure 2B). Significantly altered hit genes in the *SUN/koi* mutant corresponded to distinct functions. Cluster 2, predicted to include genes that were transcriptionally repressed (due to reduced Pol II binding), included genes associated with muscle function, e.g. genes coding for contractile proteins and for myosin complexes. It also included genes associated with pupal adhesion, predicted to be expressed by salivary gland cells, possibly due to leaky expression of Mef2-GAL4 in the salivary gland. Cluster 1, predicted to include genes that were also transcriptionally repressed due to increased binding of Polycomb and HP1a, contained genes coding for components of mTOR signaling, involved in muscle cell growth (Yoon, 2017). Repression of mTOR signaling might result in inhibition of muscle growth that was indeed observed in the SUN/*koi* mutant muscles. Figure 2C includes a list of specific genes of interest, and their fold change values. It includes the identity of 26 muscle-specific genes mostly predicted to be transcriptionally down regulated in the *SUN/koi* mutant muscles (either through decreased RNA-Pol II or increased Polycomb binding). Such transcriptional repression explains the significantly thinner muscle fibers and impaired contraction of the *SUN/koi* mutant muscles. Interestingly we also observed decreased binding of RNA-Pol II and increased binding of HP1a to a group of genes involved in nuclear transport, predicting impaired functionality of the muscle fibers as well. In addition, a group of transcriptional regulators showed increased Polycomb binding and decreased RNA-Pol II binding, again predicting impaired function of the muscle fibers, and explaining the weakness of the larval muscles and their thinner phenotype, resulting in slower larval movement.

**Figure 2.**
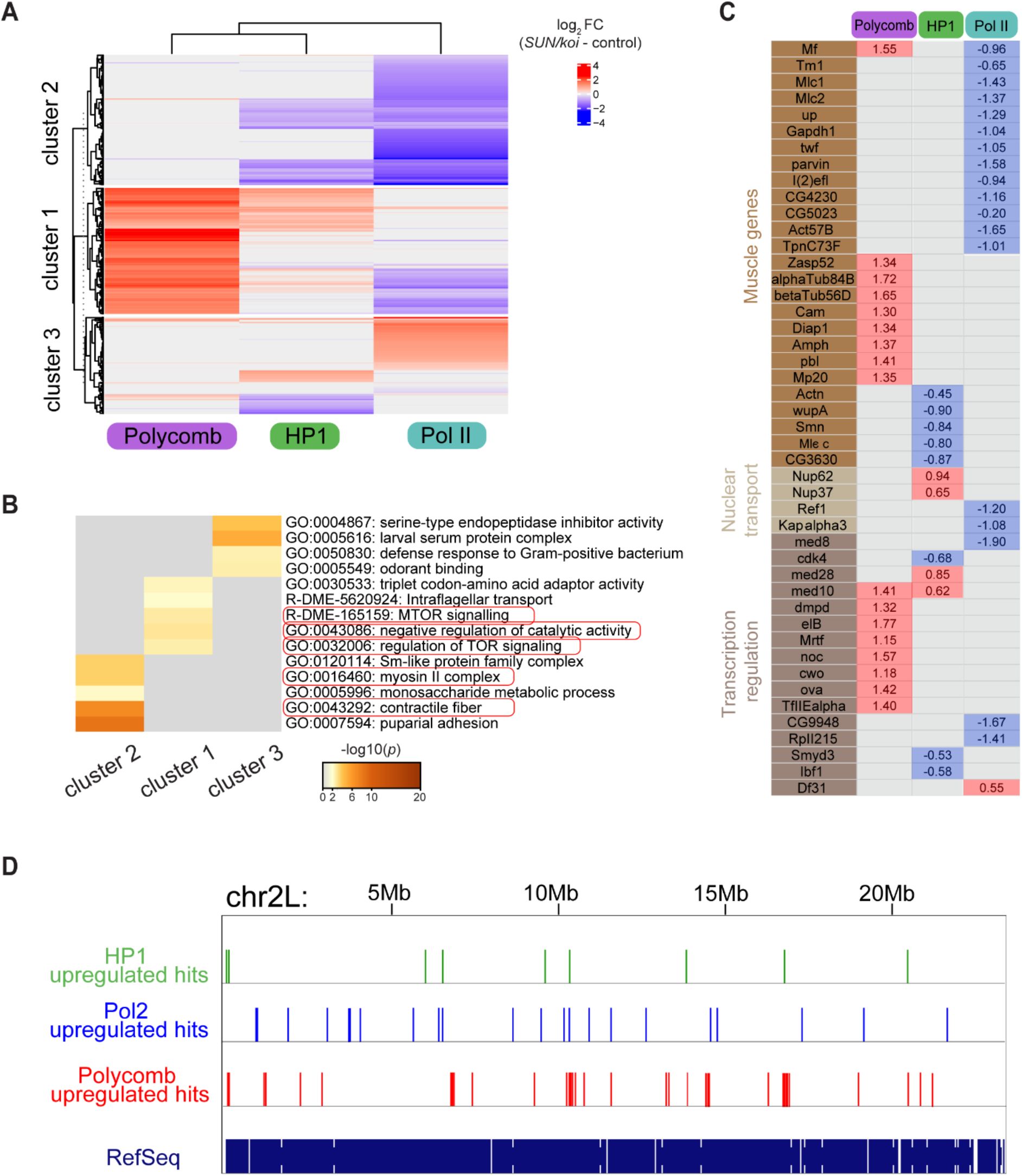
Polycomb, HP1a, and RNA-Pol II binding regulate distinct group of genes in the LINC mutated muscle fibers, with overall increased Polycomb repression and decreased RNA-Pol II activation. **(A)** The difference in *SUN/koi* and control fold change occupancy represented as heatmap for significantly altered genes in the Polycomb, HP1a, and RNA-Pol II groups. Non-significant genes are labeled in gray. K-means gene clustering reveals down-regulated genes in cluster 1 (increased Polycomb binding), and cluster 2 (decreased RNA-Pol II binding), and heterogenous genes in cluster 3. **(B)** GO enrichment analysis (heatmap of *p*-values) on the three clusters of genes identified in (A). **(C)** The difference in *SUN/koi* and control fold change occupancy for specific genes of interest. (D) Representative image of the distribution of the upregulated HP1a, RNA-Pol II, and Polycomb hits on chromosome 2L from DamID analysis, indicating increased clustering of Polycomb upregulated genes along the genome.

Taken together our genome-wide analysis indicates that defective LINC complex associates with <u>increased chromatin repression</u>, mostly through increased Polycomb binding, and decreased transcriptional activation through reduced RNA-Pol II binding. Interestingly each chromatin factor group is involved in the regulation of a distinct group of genes with only minimal overlap in genes controlled by Polycomb, HP1a, and RNA-Pol II. This is consistent with genome binding maps illustrating no overlap between Polycomb and HP1a repressive chromatin (Filion et al., 2010; De Wit et al., 2007).

To test if HP1a, RNA-Pol II, and Polycomb upregulated hits tend to cluster more than expected by chance, we run a simulation in which we randomly picked genes from the *Drosophila* dm6 genome (chr2R, chr2L, chr3R and chr3L) and counted the number of genes that are in a distance < 10,000bp (expected number of close genes). To match the HP1a, RNA-Pol II, and Polycomb upregulated hits distributions over the genome, we picked the same number of genes, corresponding to the upregulated positive hits for each group. The process was repeated Nruns=1000 times and the p-values were then calculated for the fraction NGE+1/Nruns+1, where NGE is the number of simulations with a value greater than or equal to the number of hits for each factor. This analysis indicates that the Polycomb upregulated genes tend to cluster along the chromosomes significantly more compared to randomly distributed genes, namely, 14 Polycomb upregulated genes were located within 10kbp compared to a median of 7 from Monte-Carlo simulation (p=0.012, number of gene hits n=136) (Figure 2D). The number of upregulated gene hits for RNA-Pol II and HP1a was significantly lower, however a similar tendency for clustering was observed, where 9 RNA Pol II upregulated genes were located within 10kbp compared to a median of 5 from Monte-Carlo simulation (p=0.002, and n=75) and 9 HP1a upregulated genes clustered within 10bp compared to a median of 5 from Monte-Carlo simulation (p=0.001, n=65). This analysis indicated a change in chromatin distribution in the *SUN/koi* mutants where Polycomb hits tend to cluster along the chromosome’s length.

### Perturbed LINC complex associates with increased epigenetic chromatin repression, and reduced epigenetic chromatin activation in muscle nuclei

The changes in Polycomb and RNA-Pol II genome binding led us to address possible changes in the meso-scale epigenetic landscape in *SUN/koi* mutated muscle fibers of 3^rd^ instar larvae, using antibody staining for three epigenetic modifications, namely H3K9ac, H3K27me3, and H3K9me3. Figure 3A shows representative muscle nuclei labeled with the repressive chromatin modifications H3K27me3 (purple), and H3K9me3 (green), with overall increased signal of both marks, in the *SUN/koi* mutated muscle nuclei. Figure 3B shows quantification for the mean nuclear fluorescence intensity with 43% increase in H3K27me3 (p<0.05), and 82% increase in H3K9me3 (p<0.01). The overall increased intensity of repressive chromatin marks is in agreement with the increased Polycomb, and HP1a binding observed with the targeted DamID. As previously reported, the *SUN/koi* mutant displays smaller and variable myonuclear size and increased DNA ploidy (Wang et al., 2018). We therefore asked whether the observed epigenetic changes correlate with the nuclear size. Figure 3C shows the mean H3K27me3, and H3K9me3 fluorescence intensity, for each nucleus, as function of the nuclear volume (log10 scale). Interestingly, both of the repressive marks displayed increased repression with smaller nuclear volumes. We performed linear mixed model analysis on the dependence of mean H3K27me3 intensity (left), and mean H3K9me3 intensity (right) on nuclear volume in the *SUN/koi* group, compared to control (red and black dots, respectively). We first compared fitted mixed models for all pulled nuclear volumes and found significant differences between the *SUN/koi* and control groups, for H3K27me3 and H3K9me3 repressive marks (p<0.01). To assess the contribution of the smaller nuclei, which are present only in the *SUN/koi* groups, we then fitted mixed models only for the larger nuclei, with overlapping volumes between the *SUN/koi* and control groups. The larger, overlapping volume nuclei showed no significant difference between the *SUN/koi* and control models, for both repressive marks (p>0.2), suggesting that the smaller nuclei in the *SUN/koi* groups are the major contributors to the increased H3K27me3 and H3K9me3 repression. To address whether the increased repressive epigenetic modifications represent muscles lacking LINC function we characterized larval mutant muscles deficient of both *Msp300* and *klar* using a double mutant combination of *klar;Msp300* lacking the KASH domain (ΔKASH) (Xie and Fischer, 2008). We found a similar tendency in the ΔKASH mutants (Supplemental Figure S1 A-B), with 48% increase in mean fluorescent H3K27me3 intensity (p<0.001) and 25% increase in mean fluorescent H3K9me3 intensity (p<0.001) signals, and the repressive mark intensity inversely correlated with nuclear volume (Supplemental Figure S1 C-D).

**Figure 3.**
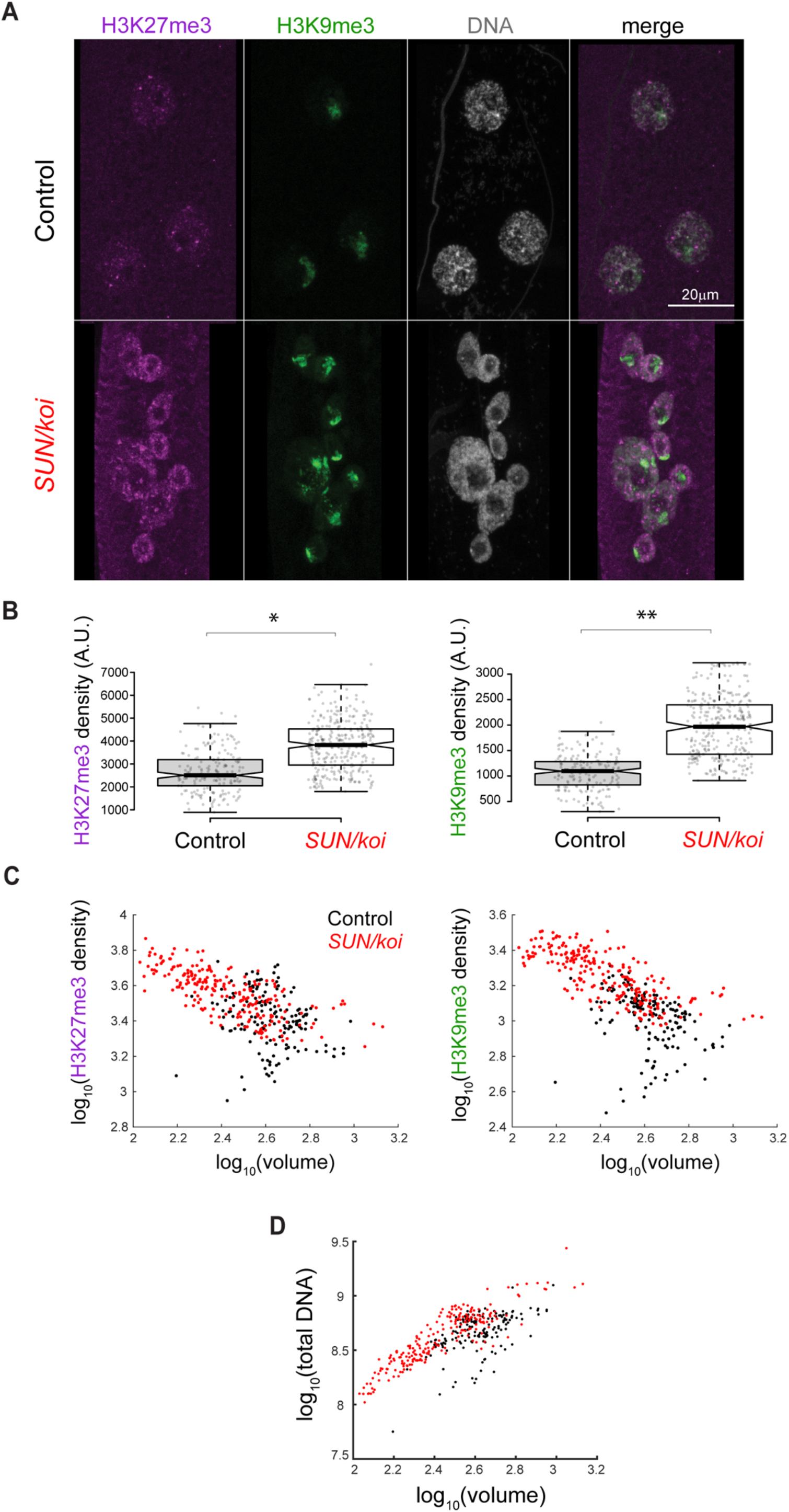
Increased repressive H3K27me3 and H3K9me3 chromatin density, inversely correlated with nuclear volume, in *SUN/koi* mutated muscle fibers. **(A)** Muscle nuclei labeled with H3K27me3 (purple), H3K9me3 (green), and Hoechst for total DNA (gray). **(B)** Quantification of mean nuclear fluorescence intensity shows increased H3K27me3 and H3K9me3 in *SUN/koi* mutated muscle fibers. **(C)** Mean nuclear fluorescence intensity is plotted against the corresponding nuclear volume (log10 scale) for each epigenetic mark. Significant difference in the linear mixed model fit between the *SUN/koi* and control groups, for both repressive marks (p<0.01). Similar analysis comparing only the larger nuclei (overlapping in *SUN/koi* and control groups) shows no significant difference between the fits, suggesting that mostly the smaller nuclei in the *SUN/koi* mutant contribute to the increased H3K27me3 and H3K9me3 repression. N=5 larvae, n=177 nuclei in control, N=5 larvae, n=295 nuclei in *SUN/koi*. For statistical significance *p<0.05, **p<0.01. (D) Total DNA intensity plotted versus nuclear volume (log10 scale) for *SUN/koi* (red) and control (black). Linear mixed model analysis confirms significant difference between the *SUN/koi* and control fits (p<0.01), with left-ward shift in the total DNA – nuclear volume relationship for the *SUN/koi* group. N=5 larvae, n=177 nuclei in control, N=5 larvae, n=295 nuclei in *SUN/koi*.

*Drosophila* muscle nuclei are polyploid, and we showed previously that chromatin volume scales linearly with nuclear volume, maintaining chromatin volume fraction constant (Amiad-Pavlov et al., 2021). The inverse correlation of chromatin repression with nuclear volume, observed in the *SUN/koi* mutant muscles, suggested increased chromatin condensation which might contribute to the excessive chromatin repression (Schuettengruber et al., 2017). To address this, we compared the global (nuclear scale) DNA condensation in *SUN/koi* and control myonuclei. Figure 3D shows a plot of the total DNA intensity as a function of nuclear volume (log10 scale) for *SUN/koi* and control groups (red and black dots, respectively). A linear mixed model analysis confirms a significant difference between the *SUN/koi* and control fits (p<0.01) with a leftward-shift in the *SUN/koi* total DNA-volume relationship, indicating that for each *SUN/koi* nucleus the DNA is condensed in a smaller nuclear volume, relative to control. Interestingly, we did not observe similar trend of DNA condensation in the ΔKASH mutant muscle fibers (Supplemental Figure S3).

### Increased H3K27me3 repression is specific for mature LINC mutant muscles

We further asked whether the increased H3K27me3 repression in the muscles is caused by requirement for *SUN/koi* in the course of muscle differentiation during embryonic stages, or is it observed also in fully differentiated muscle fibers. For that we conditionally knocked-down *SUN/koi* by RNAi in larval muscle fibers, after muscle differentiation has been fully completed, utilizing the temperature sensitive *GAL80/Mef2-GAL4* driver. Whereas muscle specific *SUN/koi* knockdown induced after muscle differentiation (in hatched larvae) did not cause significant defects in nuclear positioning, nor did it affect nuclear volume, it did recapitulate the increased levels of H3K27me3 modification with 45% increase in the mean nuclear fluorescent intensity (Figure 4A-B, p=0.011). Consistently, the temporal *SUN/koi* knockdown in differentiated muscles showed no dependance of H3K27me3 levels on nuclear volume (Figure 4C) and no change in the total DNA/nuclear volume relationship was observed between control and *SUN/koi*-RNAi muscle fibers (Figure 4D). These results suggest that while the *SUN/koi* is required to maintain nuclear volume, optimal DNA density, and nuclear positioning during embryonic development, proper LINC function in mature muscles is required to inhibit epigenetic repression, independently of changes in DNA density.

**Figure 4:**
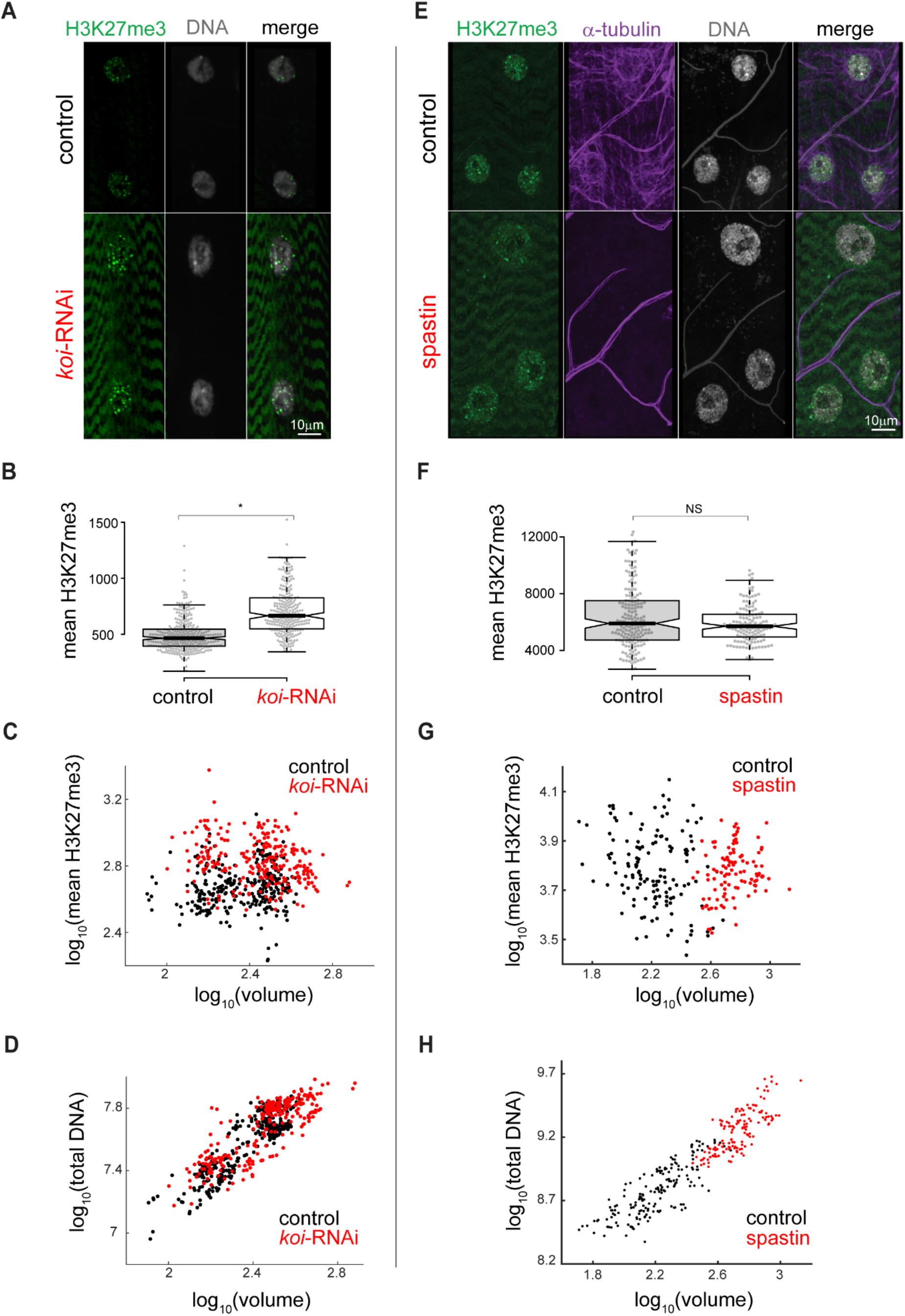
Increased H3K27me3 repression induced in temporal *SUN/koi* knockdown targeted to mature muscle fibers, but not in temporal disruption of the microtubule network. **(A)** Elevated H3K27me3 levels (green) and preserved DNA density (gray) in nuclei of mature muscle *SUN/koi*-RNAi. **(B)** Quantification of mean nuclear H3K27me3 intensity shows 45% increase in *SUN/koi*-RNAi group (p=0.011). **(C)** No correlation between mean nuclear H3K27me3 fluorescence intensity and the corresponding nuclear volume (log10 scale) in control (black) or *SUN/koi*-RNAi (red) groups **(D)** Preserved linear scaling between total DNA and nuclear volume (log10 scale) in *SUN/koi*-RNAi group. N=5 larvae, n=308 nuclei in control, N=5 larvae, n=250 nuclei in *SUN/koi*-RNAi. **(E)** Muscle nuclei of control and 1-day spastin overexpression demonstrate preserved H3K27me3 levels (green) upon MT network disruption (α-Tubulin label in purple), and preserved DNA density (gray), despite elevated nuclear volume. No change in mean H3K27me3 florescence intensity **(F)** and its scaling with nuclear volume (**G**, log10 scale) is maintained upon spastin overexpression. **(H)** Proportional increase in DNA intensity and nuclear volume (log10 scale) results in preserved global DNA density upon spastin over-expression. N=5 larvae, n=174 nuclei in control, N=5 larvae, n=140 nuclei in spastin.

Since the LINC complex is required for the cytoplasmic-nuclear force transmission, which is particularly high in mature muscle fibers, we tested whether alternative perturbation of such forces will show similar increase in repressive chromatin. This was performed by muscle specific, temporal disruption of the microtubule (MT) network, normally exerting high compressive forces on the nucleus, by overexpression of spastin, a MT severing protein, using a combination of *GAL80^ts^/Mef-GAL4* drivers with UAS-spastin. This treatment disrupted the MT in muscles for a short time period without affecting sarcomere organization or muscle size (Wang et al., 2018). Figure 4E depicts representative control and spastin overexpression muscle nuclei, with preserved H3K27me3 and DNA density upon disruption of the MT network, despite increased nuclear volume. We compared the H3K27 tri-methylated fluorescent signal, and demonstrate that the mean fluorescent intensity of H3K27 tri-methylation (Figure 4F), as well as the ratio between mean H3K27 tri-methylation fluorescent signal and nuclear volume (Figure 4G) in spastin-treated muscle nuclei did not differ from control. Figure 4H quantifies the proportional increase in nuclear volume and total DNA content with spastin overexpression (presumably as a result of DNA endoreplication), implying that DNA/nuclear volume relationships were similar to control. This result reinforces the specific and unique role of the LINC complex in inhibition of epigenetic repressive modification in the mature muscle fibers.

### Reduced epigenetic gene acetylation is observed in the *SUN/koi* mutant muscles

In agreement with the reduced RNA-Pol II binding to muscle specific genes, we identify a decrease in H3K9ac chromatin modification, which marks the active promotors, in the *SUN/koi* mutant muscle fibers (Figure 5A). Figure 5B quantifies the mean nuclear H3K9ac intensity with 37% reduction in the *SUN/koi* mutant (p<0.01). Notably, the active H3K9ac mark showed no correlation with nuclear volume (Figure 5C). Interestingly, the ΔKASH mutants did not show similar tendency as the *SUN/koi* mutants, as we observed no significant change in the mean nuclear H3K9ac intensity (p=0.19) (Supplemental Figure S2). Overall, this data indicate reduced active landscape in the *SUN/koi* that is consistent with the reduced binding of RNA Pol II to an array of genes, including muscle genes.

**Figure 5.**
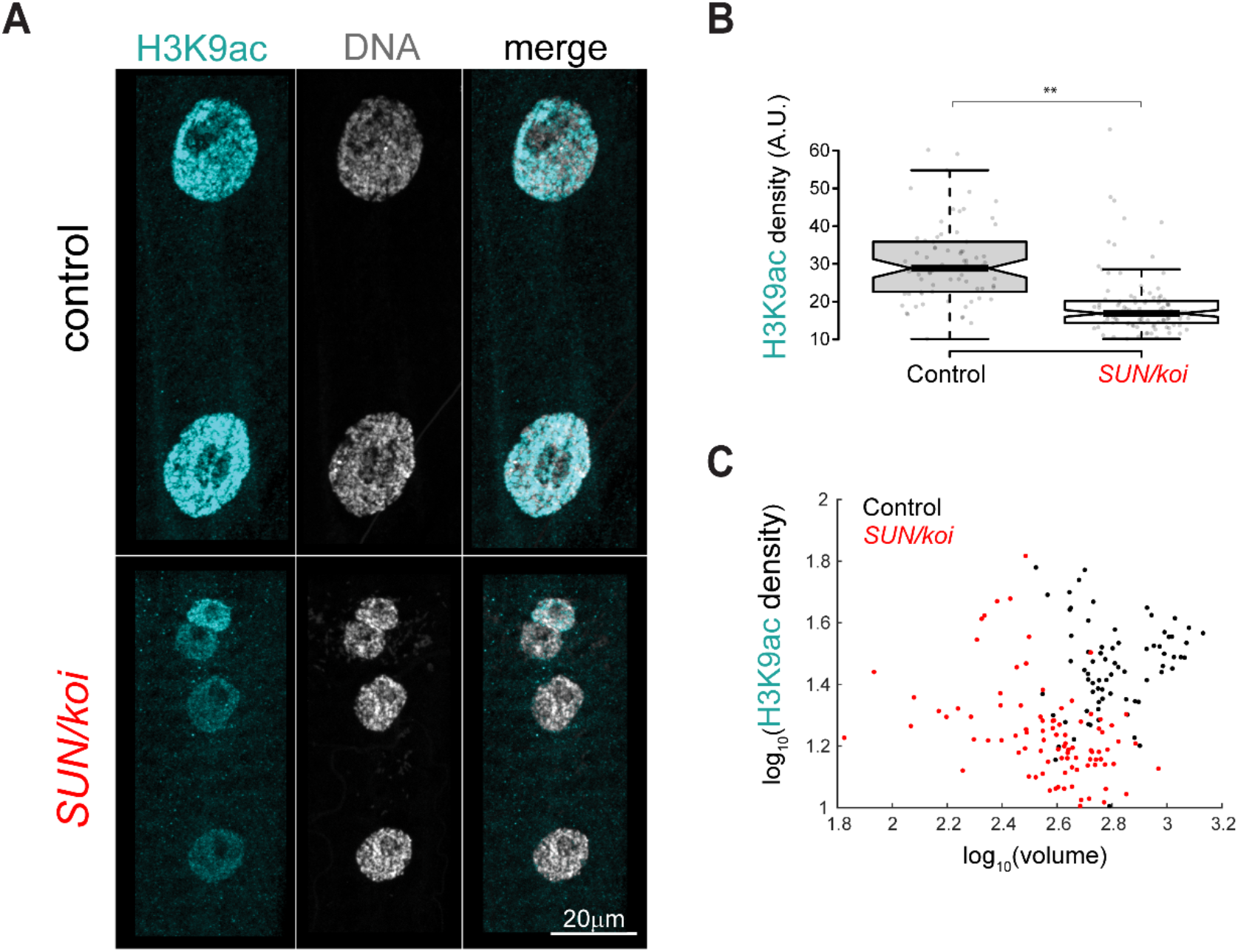
Reduced active H3K9ac chromatin density in *SUN/koi* mutated muscle fibers, independent of nuclear volume. **(A)** Muscle nuclei labeled with H3K9ac (cyan) and Hoechst for total DNA (gray). **(B)** Quantification of mean nuclear fluorescence intensity shows decreased H3K9ac in *SUN/koi* mutated muscle fibers. **(C)** Mean nuclear H3K9ac intensity is plotted against the corresponding nuclear volume (log10 scale) and shows no significant correlation with nuclear volume between the control and *SUN/koi* groups (left). N=5 larvae, n=73 in control, N=4 larvae, n=103 nuclei in *SUN/koi*. **p<0.01.

### Live imaging of the 3D distribution of H3K27me3 sites reveals increased cluster size in *SUN/koi* mutated muscle fibers

To investigate the spatial distribution of H3K27me3 sites in the *SUN/koi* mutant muscle nuclei we performed live 3D imaging of larval muscles utilizing the H3K27me3-GFP mintbody (Tjalsma et al., 2021). Representative 3D spatial distribution of H3K27me3-GFP in control and *SUN/koi* mutated nuclei is shown in Supplemental Movie S1 and Supplemental Movie S2, respectively. Figure 6 shows middle confocal sections of representative control (left) and *SUN/koi* mutant (right) muscle nuclei labeled with H3K27me-GFP mintbody. The strong H3K27me3 signal appears punctuated within the nucleus, with weaker nuclear and cytoplasmic background. Compared to control distribution, the H3K27me3 puncta in the *SUN/koi* mutant nuclei appeared more spread-out throughout the chromatin, resulting in larger repressive H3K27me3 hubs. We quantified the spread of the puncta by measuring the volume (Figure 6B) and the number (Figure 6C) of strong H3K27me3 puncta in each group. The *SUN/koi* puncta volume is on average 149% higher than control (p<0.01) with no significant change in the number of the puncta between the groups. Thus, the live 3D distribution of H3K27me3 puncta demonstrates increased spreading of this repressive mark to neighboring chromatin regions in the *SUN/koi* mutated muscle nuclei, further supporting the clustering of positive Polycomb binding location across the genome (Figure 2D). Considering the larger H3K27me3 puncta in live *SUN/koi* mutant muscle nuclei, together with the increased Polycomb binding (Figure 1A) and the clustering of positive Polycomb binding sites (Figure 2D), our findings suggest that *SUN/koi* disruption induces increased repression by the spread of repressive H3K27me3 chromatin modifications to nearby chromatin regions.

**Figure 6:**
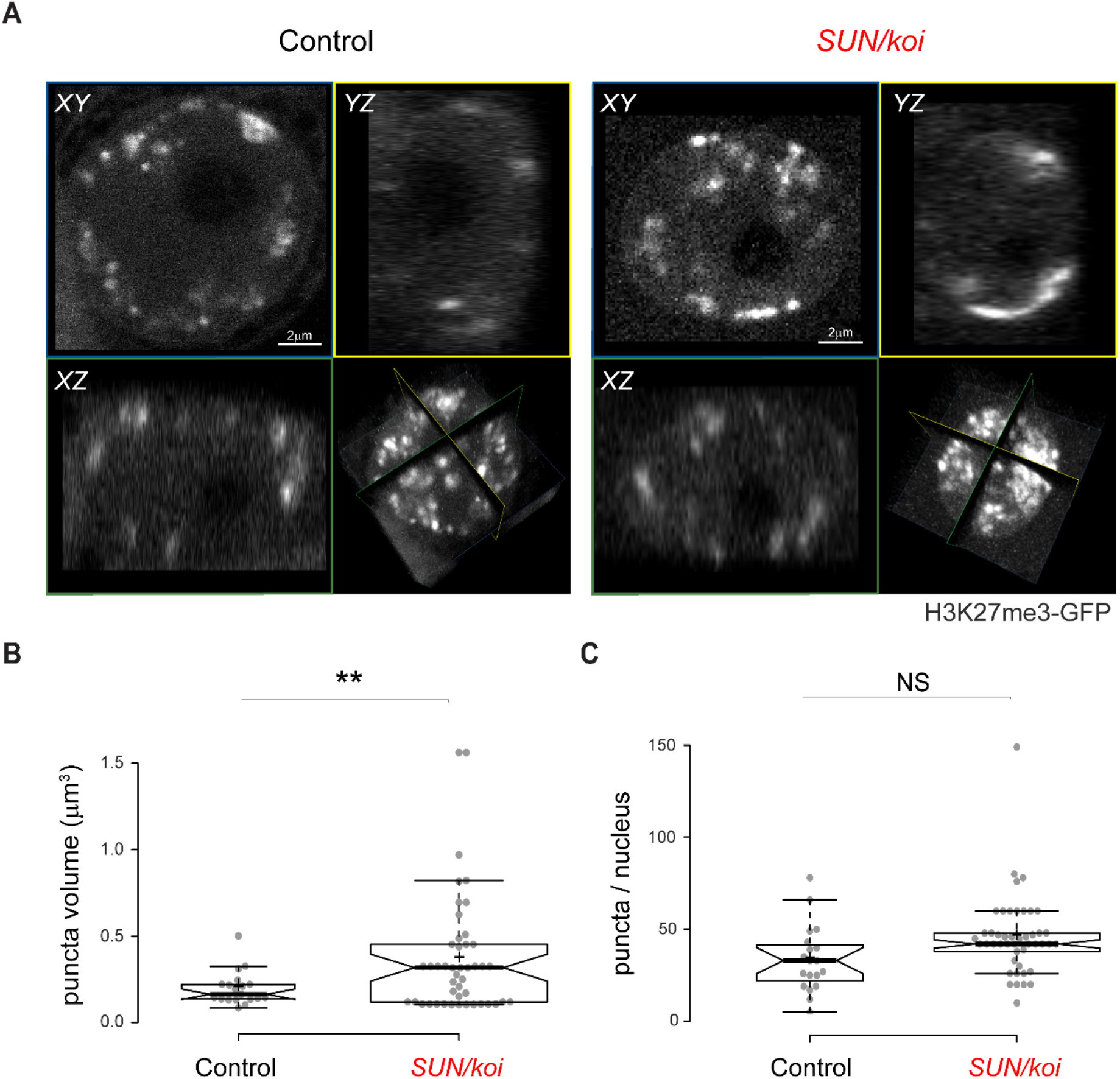
Live 3D imaging of H3K27me3 mintbody distribution indicates increased spread of the repressive mark in *SUN/koi* mutant muscle nuclei. **(A)** Middle confocal planes of control (left) and *SUN/koi* (right) muscle nuclei show punctuated repressive regions throughout the nucleus. Increased mean puncta volume (B) and unchanged number of puncta per nucleus (C) in the *SUN/koi* mutant group. N=4 larvae, n=19 nuclei in control, N=4 larvae, n=46 nuclei in *SUN/koi* (**p<0.01).

### Dissociation of H3K27me3 chromatin from the lamina is linked to increased H3K27me3 clustering in LINC mutant muscle

Our previous work indicated peripheral chromatin organization in nuclei of mature muscle fibers, which is sensitive to lamin A/C levels, when imaged live, *in-vivo* (Amiad-Pavlov et al., 2021). To assess the meso-scale chromatin organization in *SUN/koi* mutated muscles we imaged live larval myonuclei co-labeled with H2B-mRFP and Klar-GFP. However, in the *SUN/koi* mutant, the Klar-GFP did not localize to the nuclear envelope, due to the lack of *SUN/koi* (Figure S4). Nevertheless, we analyzed the live, 3D distribution of chromatin by quantifying H2A-GFP density in radial shells from the nuclear periphery to the nuclear center (Figure 7A). *SUN/koi* mutated muscle nuclei exhibit altered chromatin distribution with reduction in peripheral chromatin density and a shift towards the center.

**Figure 7:**
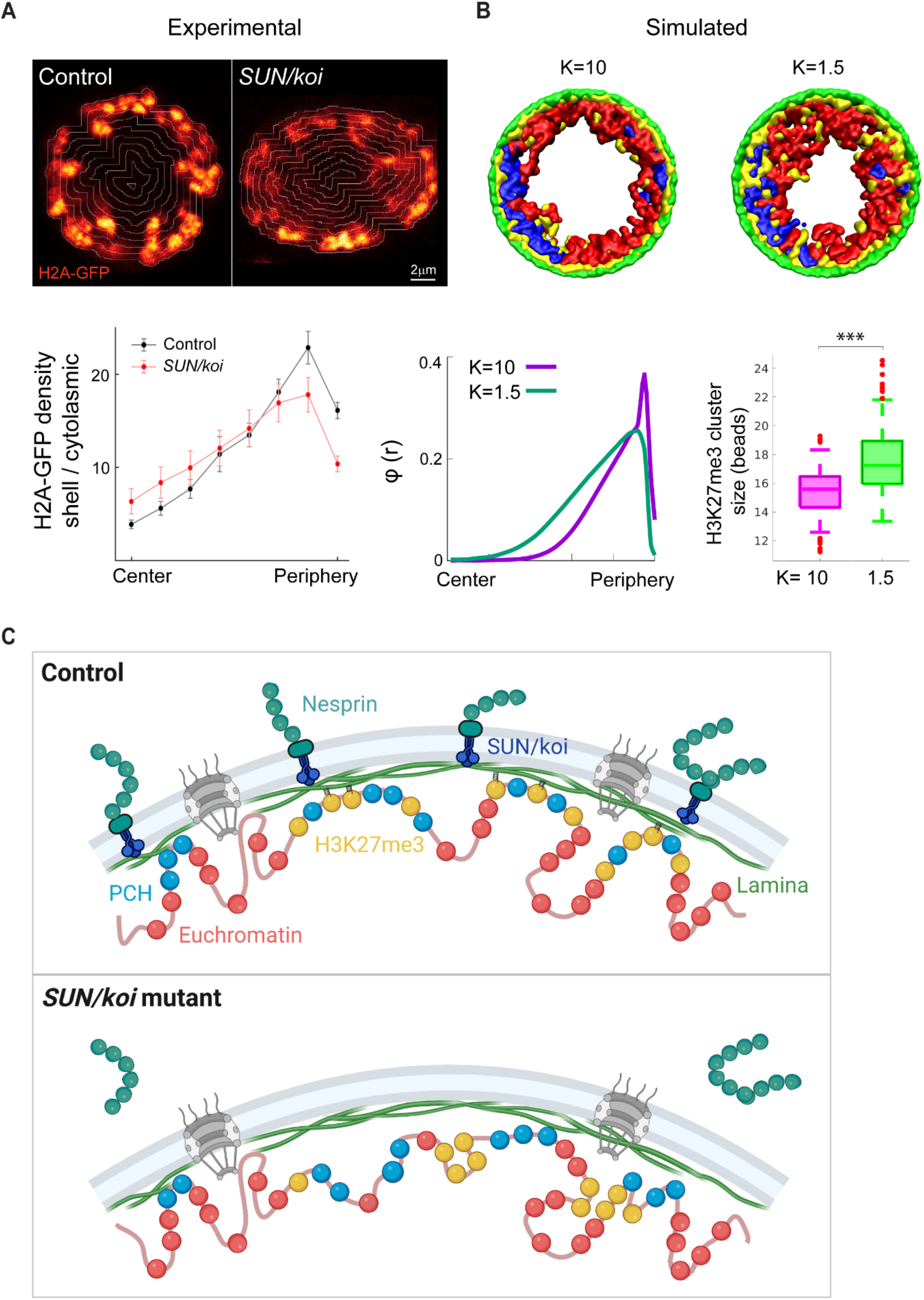
Experimental and computational evidence for chromatin redistribution towards the center as the mechanism for increased repressive clustering in the LINC mutant. (A) Radial chromatin distribution analysis from live, 3D imaging. Middle confocal planes of control (left) and *SUN/koi* (right) muscle nuclei labeled with H2A-GFP. Mean chromatin density is quantified for the radial shells and demonstrate decreased peripheral chromatin density with a central shift in the *SUN/koi* mutants (red, N=4 larvae, n=11 nuclei), compared to control (black, N=3 larvae, n=10 nuclei). (B) Computational model to describe the *Drosophila* chromosomes with a flexible bead-spring polymer chain containing three distinct regions: euchromatin (red), pericentromeric heterochromatin (blue), and H3K27me3 modification (yellow). The polymers are confined by a spherical wall comprising immobile, lamin beads (green). The yellow (H3K27me3) beads are attracted to the green (laminar) beads and their bonding strength is modeled by a dynamic, harmonic (spring-like) potential K. Simulation snapshots (XY plane view), and quantification of the radial chromatin bead density demonstrate that for strong bonding of the yellow (H3K27me3) and green (laminar) beads, K=10, most of the yellow clusters are located near the lamina, while for weak bonding, K=1.5, the yellow clusters diffuse within the spherical volume, resulting in central chromatin shift. Box plots show increased H3K27me3 bead cluster size with decreased bonding strength K (p<0.001). (C) Schematic model to represent the mechanism for LINC mediated inhibition of chromatin repression. *SUN/koi* mutation decreases repressive chromatin binding to the lamina, resulting in increased H3K27me3 clustering in the nucleoplasm and redistribution of chromatin inward from the periphery (created with BioRender.com).

Experimentally, the live 3D analysis in this study suggests that disruption of the LINC complex, results with excessive H3K27 tri-methylation, chromatin redistribution from the periphery towards the center of the nucleus, and extra Polycomb binding to genes hubs, presumably leading to increased size of repressive H3K27me3 clusters. We therefore studied a computational model to explore a possible link between dissociation of the chromatin, including tri-methylated H3K27 hubs, from the nuclear envelope and the formation of larger repressive clusters in LINC mutants. Toward this end, we introduced a coarse-grained model for the *Drosophila* genome that incorporates binding/unbinding of H3K27me3-modified chromatin (facultative heterochromatin) to the lamina. The *Drosophila* genome is simulated by beads connected by springs to model each chromosome, where a bead of diameter *σ* = 30 nm includes 5 kbps of DNA. We confine four such flexible polymers, which represent the chromosomes of *Drosophila*, to a sphere which represents the nucleus. The chromosomal chains consist of three types of beads: euchromatin beads, pericentromeric heterochromatin (PCH) beads, and H3K27me3 modified beads (Figure 7B, red, blue, and yellow beads respectively). In our model, euchromatin and heterochromatin (PCH and H3K27me3) are distinguished by *ε* -the strength of their Lennard-Jones attraction (LJ, see further details in the methods section). H3K27me3 beads form bonds with nearby lamina beads with a fluctuating H3K27me3-lamin bead distance after bond formation. The change in the bond energy as a function of the H3K27me3-lamin bead separation is described by a harmonic (spring-like) potential with a spring constant K. Our simulation results show that decreasing the spring constant K, namely the H3K27me3-lamina bond strength, increases the average H3K27me3 cluster size in the nucleoplasm, resulting in a chromatin density shift from peripheral towards more central (Figure 7B). This occurs because smaller values of K lead to increased probability of H3K27me3-lamin unbinding. The H3K27me3 that has unbonded from the lamin, phase separates in the nucleoplasm due to the attractive interactions of the H3K27me3 beads; this increases chromatin volume in the nucleoplasm relative to periphery. Overall, our experimental and computational results suggest a model whereby disruption of the LINC complex in the mature muscle weaken the interactions of repressive chromatin regions with the lamina, resulting in increased repressive H2K27me3 clustering in the nucleoplasm, and overall chromatin shift from the periphery towards the center (Figure 7C). Functionally, the increased epigenetic repression points to the downregulation of muscle contractile genes in agreement with thinner muscle and perturbed mobility of the mutant larvae.

## Discussion

The LINC complex has been implicated in the regulation of chromatin 3D organization; however, its specific contribution and the underlying mechanism are yet to be elucidated. Here we addressed the contribution of the LINC complex to the regulation of chromatin organization and epigenetics. We demonstrate enhanced genome wide binding of Polycomb, altered binding profile of HP1a, and RNA-Pol II in *SUN/koi* mutant muscles, predicted to reduce transcription of genes coding for contractile proteins, as well as genes associated with muscle size. The enhanced transcriptional repression is consistent with increased levels of the repressive tri-methylated H3K27 and H3K9, and decreased levels of the active H3K9ac chromatin modifications. Whereas DNA condensation is elevated in *SUN/koi* mutant muscles due to reduced nuclear volume, we show that *SUN/koi* conditional knock-down nuclei with nuclear size comparable to control, still exhibit enhanced epigenetic repression, indicating a mechanism that is not based on DNA condensation. Remarkably, live imaging of tri-methylated H3K27 marked chromatin in *SUN/koi* mutant muscles reveals larger repressive chromatin clusters, that correlated with a transition of the chromatin from the nuclear periphery towards nuclear center. Computational modeling of the distribution of H3K27me3 sites indicates that dissociation of chromatin-nuclear lamina binding leads to the formation of larger H3K27me3 repressive clusters, consistent with our observations. Mechanistically we suggest that weakening of chromatin binding to the nuclear lamina in the *SUN/koi* mutants, leads to excessive clustering of H3K27me3 sites and to spreading of Polycomb binding to neighboring genes. Taken together these results suggest that the LINC complex is required for inhibition of excessive chromatin epigenetic repression necessary for robust transcriptional regulation in fully differentiated muscle fibers.

Polycomb genome-wide binding profile showed a clearly directed trend with increased binding to a group of 143 genes (out of total 148 genes with altered binding) observed in the *SUN/koi* mutant muscles, which implies a significantly enhanced transcription repression. Interestingly, among these genes (cluster 1) we identified enrichment for two GO terms functionally associated with TOR signaling, a major signaling pathway regulating muscle growth. Enhanced transcriptional repression of TOR signaling in the *SUN/koi* mutants is consistent with the thinner muscle phenotype observed in these muscles (Elhanany-Tamir et al., 2012; Wang et al., 2018). Furthermore, since Polycomb binds specifically to tri-methylated H3K27-marked chromatin, the enhanced Polycomb binding is also consistent with an increased H3K27me3 fluorescent labeling observed in the *SUN/koi* mutant muscles. At this stage we cannot determine whether increased polycomb binding leads to increased H3K27me3 or vice versa, since Polycomb was demonstrated to self-aggregate with chromatin due to phase separation (Tatavosian et al., 2019), but also, to further promote tri-methylation of H3K27 (Schuettengruber et al., 2017). Our observation that in muscles of live *SUN/koi* mutant larvae the H3K27me3-positive puncta grew in size is consistent with increased self-aggregation of pre-existing H3K27me3-associated chromatin clusters.

Polycomb is a member of polycomb repressive complexes (PRCs) and has been extensively studied in development and differentiating cells. PRC binding to epigenetically modified chromatin leads to transcriptionally poised state, which then prevents unscheduled differentiation (Schuettengruber et al., 2017). Therefore, enhanced binding of Polycomb to chromatin might inhibit transcriptionally regulated developmental transitions in the larval muscle fibers. The idea that such transitions might be regulated by mechanical signals, mediated by the LINC complex, is intriguing and novel. Consistent with the contribution of the LINC complex to the control of developmental transitions is a recent description of differentiating keratinocytes deficient of both SUN1 and SUN2, which undergo precocious differentiation transition (Carley et al., 2021). In contrast to our results, Carley et al. demonstrated that lack of SUN proteins led to reduced repression, higher accessibility of the chromatin, and activation of terminal keratinocyte differentiation. However, this difference might stem from a distinct state of cellular differentiation of each experimental system studied. Whereas our analysis was performed on fully differentiated muscle fibers, the analysis of SUN-deficient keratinocytes was performed on keratinocytes prior to their terminal differentiation. Since chromatin mesoscale viscoelastic properties change during cell differentiation (Matsushita et al., 2021), the different outcome might result from the differences in chromatin properties in undifferentiated versus differentiated cells. Importantly however, the LINC complex appears to be essential for preservation of the epigenetic state of the chromatin in both cases.

A number of studies suggested a link between nuclear mechanical stimulation and chromatin epigenetic repression. For example, long-term stretch of epidermal progenitor cells led to increased H3K27me3 occupancy resulting with silencing of epidermal differentiation (Heo et al., 2016; Le et al., 2016). Furthermore, mechanical inputs inducing nuclear deformation led to changes in the distribution of heterochromatin and euchromatin in various cell types, including cardiomyocytes and keratinocytes (Seelbinder et al., 2021; Wagh et al., 2021). Mechanistically it has been proposed that nuclear deformation changes the extent of chromatin binding to the nuclear envelope, affecting primarily repressed chromatin (Wagh et al., 2021). Interestingly, epidermal progenitor cells subjected to mechanical stretch exhibited reduced H3K9 tri-methylation associated with HP1a binding, accompanied by concomitant increase of Polycomb repressive complex 2 (PRC2) binding, resulting with enhanced global repression (Le et al., 2016). Our analysis of LINC mutant muscles, with perturbed mechanical signal transduction into the nucleus, did not reveal similar replacement between the binding of Polycomb and that of HP1a, and suggests that each of these transcription repressors changed its binding independently of the other on mostly distinct set of genes, while both H3K9me3 and H3K27me3 repressive marks were significantly increased in the LINC mutant muscles.

RNA-Pol II binding profile in *SUN/koi* mutant muscles reveals genes in cluster 2 with decreased binding to the transcriptional machinery complex. According to GO analysis this cluster contains a group of genes coding for muscle contractile proteins. Reduced transcription of such genes is expected to result with muscle weakening and failure in larval locomotion and indeed both phenotypes were observed in the *SUN/koi* mutants. Noteworthy, because the DamId analysis was performed at late stage of 3^rd^ instar larvae, some of the muscle-specific genes undergo transcription reduction due to transition from larvae to pupal stage (flybase.org). For example, muscle specific genes including myosin heavy chain (MHC), *sallimus (Drosophila* titin-like gene), *MSP300 (*Nesprin-like gene), *alpha actinin (Actn)* and others show relative reduced mRNA levels at late 3^rd^ instar larval stage. In such genes alterations of RNA-Pol II binding in the mutant muscles might not be detectable. Nevertheless, a group of other muscle genes did show a significantly reduced binding to RNA-Pol II e.g. *myofilin* (*Mf*), *tropomyosin1* (*Tm1)*, *myosin light chain* (Mlc), *actin 57B* (Act57B), *troponinC73F* (TpnC73F) and others. In addition, a distinct group of muscle genes showed increased Polycomb binding (e.g., *Zasp52, Amph, betaTub56D* and others), predicting their increased repression in the *SUN/koi* mutant muscles. Lastly, transcription itself might be attenuated due to the increased binding of Polycomb and HP1a to components of the transcription machinery such as mediator complex proteins *med10* and *med28*. The decreased levels of H3K9 acetylation in the *SUN/koi* mutant muscles is consistent with the paralleled reduction in RNA-Pol II binding to genes in cluster 2. Importantly, the lack of overlap between the group of genes that showed increased Polycomb binding (cluster 1) and the group of genes that exhibited decreased RNA Pol II binding (cluster 2) suggests that although both processes are regulated by the LINC complex, they might result from distinct effects of this complex on repressed and active chromatin regions.

Although enhanced DNA condensation and compaction could explain the upregulation of epigenetic repression in the LINC mutant nuclei, we do not favor this explanation since we found that elevated epigenetic repression is observed also in nuclei of normal size with conditional *SUN/koi* knock-down. The simulation results described in Figure 7B provide an alternative mechanism whereby repressive H3K27me3 clusters grow in size if chromatin affinity to the nuclear lamina is reduced, compared to the wild type situation. Experimentally we show that chromatin distribution indeed shifts from the nuclear periphery towards the nuclear center in the *SUN/koi* mutant nuclei, implicating weakening of chromatin - nuclear lamina binding. Consistently, recent studies indicated that in LINC mutants the LEM domain protein Emerin, which together with BAF promotes chromatin tethering to the lamina, fails to oligomerize at the inner nuclear membrane (Fernandez et al., 2022), and consistently BAF is eliminated from the nuclear envelope (Unnikannan et al., 2020); both evidence predict chromatin dissociation from the nuclear envelope in the LINC mutants as was observed. Therefore, we favor a mechanism by which the LINC complex stabilizes chromatin association with the nuclear envelope, preventing excessive epigenetic repression.

In summary, this study analyzed the contribution of the LINC complex to chromatin epigenetics and the binding of transcriptional regulators in fully differentiated muscle fibers. We demonstrate that in the absence of functional LINC complex chromatin epigenetic repression is enhanced, mainly due to enhanced binding of Polycomb and excessive clustering of repressed chromatin, while Pol II binding to muscle specific genes decreases. This forms the basis for abrogated muscle structure and function in LINC mutant observed in striated muscles, as well as in LINC associated cardiomyopathies.

## Materials and Methods

### Fly stocks and husbandry

The following stocks were used: koi^84^/Cyo-dfd-eYfp (FBst0025105) have been described previously (Kracklauer et al., 2007), koi-RNAi (FBst0040924), delta KASH (MSP-300^deltaKASH^;klar^mCD4^/TM6B from J.A Fischer, University of Texas, Austin, TX), tubP-GAL80^ts^/TM2 (FBst0007017), GAL4-Mef2.R (FBst0027390),, UAS-Spastin (obtained from V. Brodu, Institute Jacques Monod, Paris), ubi-H2B-mRFP/CyO; Dr/TM6B (Bosveld et al., 2017), UAS-klar-GFP (Elhanany-Tamir et al., 2012), Tub-Gal80ts; Mef2-Gal4 (obtained from F. Schnorrer IBDM, Marseille), UAS-2E12LI-EGFP(III)/TM6B (live H3K27me3-GFP mintbody obtained from H. Kimura, Tokyo Institute of Technology), His2Av-GFP (FBst0005941). Fly lines for Targeted Dam-ID (obtained from A. Brand, The Gurdon Institute, University of Cambridge): UAS-LT3-Dm (FBtp0095492), UAS-LT3-Dm-RpII215 (FBtp0095495), UAS-LT3-Dam-Pc, UAS-LT3-Dam-HP1a (Marshall and Brand, 2017).

All crosses were carried and maintained at 25°, 18°, or 29°C and raised on cornmeal agar. Homozygous *SUN/koi* and ΔKASH mutant larvae were selected by non-Cyo-YFP, and confirmed with mis-localization / aggregation phenotype of muscle nuclei. Temporal *SUN/koi* knockdown (*koi*-RNAi) and overexpression of spastin was performed using a combination of Mef2Gal4 and tubGal80ts drivers as follows: embryo collection was performed at 25°C for 12h, followed by transfer to a permissive temperature of 18°C. Larvae were transferred to the restrictive temperature of 29°C for 3 days (*koi*-RNAi) or 1 day (spastin) before late 3^rd^ instar (wandering larvae). Tub-Gal80ts; Mef2-Gal4 fly line was used as control for *SUN*/*koi* mutant, *koi*-RNAi and spastin over-expression experiments.

### Targeted DamID

We drove the expression of Dam-Polycomb, Dam-HP1a and Dam-Pol II and Dam only in control and *SUN/koi* mutant muscles using Mef2-gal4 driver (total 8 genotypes). Temporal control of Dam expression was achieved with a ubiquitously expressed temperature sensitive gal80. Each genotype contained 3 independent replicates, with 25 larvae per group. Eggs were laid for 6 hours and developed in 18°C until late larval second instar stage, and transferred to 29°C for 10 hours to allow muscle specific Dam expression. Larvae were then dissected and stored at −80°C until all triplicates were further processed together following previously described protocol (Marshall et al., 2016). Briefly, genomic DNA was extracted and digested with methylation-specific DpnI enzymes that cleave at GATC sites. A double-stranded oligonucleotide adaptor was used to ensure directional ligation. The ligation was followed by digestion with DpnII that cuts only unmethylated GATCs. Finally, a PCR primer was used to amplify adaptor-ligated sequences. These amplified sequences were deep sequenced, the results were further analyzed by a bioinformatic pipeline (Marshall and Brand, 2015), and reads were normalized to filter out nonspecific Dam binding.

Each sample was processed separately by the damid-pipeline to generate normalized ratios for each pair of samples; Dam fused to protein of interest versus Dam only (https://github.com/owenjm/damidseq_pipeline). Mean occupancy per gene was then calculated using polii script (https://github.com/owenjm/polii.gene.call). The reference genome was taken from flybase (ftp://ftp.flybase.net/releases/FB2016_02/dmel_r6.10/) and GATC sites from the damid-pipeline website. After filtering genes with DamID false discover rate (FDR) < 0.05 (per sample), the three replicates from each group were directed to a clustering algorithm based on pairwise correlation.

To determine the genes with statistically significant occupancy in *SUN/koi* mutant, relative to control, a regression on principal component analysis was performed with a z-score for each of the points around the regression line. The cut-off criteria for significantly altered binding to a gene was set to FDR< 0.05, z-score > 1.96 (2-tailed), and GATC sites > 1, corresponding to approximately 95% confidence interval. Independently, binding profiles (log2 fold change, normalized to Dam only) of specific genes were further visualized for control and *SUN/koi* mutant using the IGV browser (Robinson et al., 2011). Gene Ontology Enrichment analysis was performed with Metascape, http://metascape.org, (Zhou et al., 2019) using gene lists from the three clusters identified by K-mean clustering, with user defined background composed of all genes identified as occupied by DamID (total 7233 genes).

### Immunofluorescence and antibodies

Quantitative immunofluorescence of epigenetics marks was performed on 3^rd^ instar, wandering larvae, that were dissected in phosphate-buffered saline (PBS) and fixed, as previously described (Wang et al., 2015). Briefly, dissected larva body walls were fixed in Paraformaldehyde (4% from 16% stock of electron microscopy grade; Electron Microscopy Sciences, 15710) for 20 min, washed several times in PBS with 0.1% TritonX-100, and mounted in Shandon Immu-Mount (Thermo Fisher Scientific). The following primary antibodies were used: rabbit anti-H3K9ac (Abcam, AB4441), rabbit anti-H3K9me3 (Abcam, AB176916), mouse anti-H3K27me3 (Abcam 6002). The following conjugated secondary antibodies were used: Alexa Fluor 555 goat anti-rabbit (Renium, #A27039) and Alexa Fluor 647 goat anti-mouse (Renium, #A21235). Hoechst 33342 (1 μg/ml; Sigma-Aldrich) was used for labeling DNA.

### Live imaging

For imaging live nuclei in their intrinsic environment, a minimal constraint device for *Drosophila* larvae was designed in our laboratory, to be placed on top of a confocal microscope stage, as previously described (Lorber et al., 2020). For stationary 3D live, in-vivo imaging of muscle nuclei, selected wandering 3^rd^ instar larva is immersed in water for ~4 hours to decrease its movement (larval movement could be restored by exposure to air). For each larva, at least three nuclei were imaged from randomly chosen muscles along the entire larval body.

### Microscopy and Image acquisition

Immunofluorescence images of epigenetic marks were acquired at 23°C on a confocal microscope Zeiss LSM 800 with a Zeiss C-Apochromat 40×/1.20 W Korr M27 lens and an Immersol W 2010 immersion medium. The samples were embedded with a coverslip high precision of 1.5H ± 5 μm (Marienfeld-Superior, Lauda-Königshofen, Germany) and acquired using Zen 2.3 software (blue edition).

Live imaging of H3K27me3-GFP was performed using an inverted Leica SP8 STED3× microscope, equipped with internal Hybrid detectors and acousto-optical tunable filter (Leica Microsystems CMS GmbH, Germany) and a white light laser excitation laser. Nuclei were imaged with a HC PL APO 86×/1.20 water STED white objective, at a scan speed of 400 Hz, a pinhole of 0.8 A.U. and bit depth was 12. Z-stacks were acquired with 0.308 μm intervals. The acquired images were visualized during experiments using LAS-X software (Leica Application Suite X, Leica Microsystems CMS GmbH).

DNA-FISH imaging was performed using a Dragonfly spinning disk confocal system (Andor Technology PLC) connected to an inverted Leica Dmi8 microscope (Leica Microsystems CMS GmbH). The signals were detected simultaneously by two sCMOS Zyla (Andor) 2048X2048 cameras, 2×2 binning, CF40 pinhole, and 12-bit depth. Images were acquired with a 63×/1.3 glycerol objective and excited with 4 different laser lines (two channels per camera): 405, 488, 561, and 637-nm laser lines.

### Image Analysis

Arivis Vision4D 3.1.2-3.4 was used for image visualization and analysis. Quantitative immunofluorescence analysis was performed with dedicated pipeline that automatically segmented multiple nuclei per stack in 3D, using denoising and Otsu threshold operation on the Hoechst channel. Nuclear volumes and total fluorescent intensities of the epigenetic marks and the DNA were calculated and exported to MATLAB R2019b (MathWorks) for further analysis.

Quantification of live H3K27me3 puncta was performed by first assessing the signal intensity profile of each nucleus and subtracting the nuclear background intensity for each nucleus individually, preserving only the bright repressive puncta. Individual puncta were then automatically segmented in 3D using Otsu operation, and puncta below volume of 0.01 μm^3^ were filtered out. Radial 3D chromatin distribution analysis was performed as previously described (Amiad-Pavlov et al., 2021) by generating consecutive radial shells within the segmented nucleus and quantifying the mean H2B-GFP intensity within the shell, divided by the mean cytoplasmic intensity.

### Computational model

The coarse-grained model simulates the 176.2 Mbps long *Drosophila* genome with N=35240 beads connected by springs to model each chromosome, where a bead of diameter *σ* = 30 nm includes 5 kbps of DNA. We confine four such flexible polymers, which represent the chromosomes of *Drosophila*, to sphere which represents the nucleus. To model the relative lengths of the euchromatin, PCH, and H3K27me3 blocks, we analyzed the experimental chip-sequence data at 5 Kbps resolution and converted it into beads unit (Li et al., 2017; Lesley Brown et al., 2018). According to this data, 70% of *Drosophila* genome regions are euchromatic and 30% are identified as PCH (Li et al., 2017). The chip-seq data of H3K27me3 markers is currently only available for euchromatin. The data indicates that 22% of the euchromatin regions (Lesley Brown et al., 2018) have H3K27me3 modifications. In addition, the data shows that the average patch length for H3K27me3 is 17 beads (or 85 Kbps), and that the variation in patch length has an exponential distribution. The H3K27me3 modifications in PCH regions are very abundant (Koryakov et al., 2011; M et al., 2022), and we assume 75-80% of the PCH is covered with H3K27me3. We therefore estimate that 40% of the *Drosophila* genome contains H3K27me3 markers based on its percentage in euchromatin and PCH. Using these values, we generated beads identified as H3K27me3 patches from an exponential distribution with a mean length of 17 beads, which covers 40% of the genome. These H3K27me3 patches were generated randomly along the genome using a Monte-Carlo method (Bajpai et al., 2021), which ensures two patches cannot overlap. Within the spherical confinement (modeled as an impenetrable well), the chromatin volume fraction is taken to be 15% (Dekker and Misteli, 2015). The nuclear lamina is modeled as a thin layer of another type of unconnected beads (lamina beads) which are localized at the surface of the spherical confinement volume. For convenience, we take the lamina beads to be fixed in position and to have the same size as the chromosomal beads. A truncated Lennard-Jones (LJ) potential (Bajpai et al., 2021) accounts for bead attractions if the distance between two beads is within 2.5*σ*, along with a short-distance, strong repulsion which prevents them from overlapping. The attraction between beads is motivated by our previous experimental observations and simulations of chromosome phase separation in live, *Drosophila* nuclei (Bajpai et al., 2021). In our model, euchromatin and heterochromatin (PCH and H3K27me3) are distinguished by the strength of their Lennard-Jones attraction (*ε*). Heterochromatin beads (*ε_HH_* = 0.5 *k_B_T*) which are more condensed in the nucleus, attract each other more strongly than euchromatin beads (*ε_Eu_* = 0.35 *k_B_T*). However, both beads are attractive since both types of chromatin are observed to be phase separated; in our model, the chains are no longer in good solvent and collapse for *ε* = 0.3 *k_B_T* (Bajpai and Safran, 2022). A bead of H3K27me3 forms a bond with a nearby lamina bead when their separation is within 1.5*σ*. Each H3K27me3 bead is restricted to form a maximum of one bond with the lamina beads. After such a bond is formed, the distance between the H3K27me3 and lamina beads can fluctuate and we use a harmonic (spring-like) potential with a spring constant K to account for the change in the bond energy as a function of the H3K27me3-lamin bead separation relative to their optimal spacing of *σ*. These bonds are broken when the fluctuating distance between beads exceeds 2.5*σ*. A detailed description of the dynamic harmonic potential (bonding/unbonding) can be found in Refs. (Bajpai et al., 2021; Adame-Arana et al., 2022). We used the LAMMPS molecular dynamics package to simulate our model system (Plimpton, 1995).

### Statistics

Quantitative immunofluorescence parameters from control and *SUN/koi* mutant nuclei were compared using a mixed linear model, with genotype as a fixed effect and with larva and muscles as random effects. The number of larvae and nuclei for each group and experiment are listed in the legends. Statistics were performed in R v.4.0.2 using the package lmerTest v.3.1-2. Linear fits to the epigenetic mark intensities, and total chromatin as function of the nuclear volume were generated using linear mixed-effects model fit. BoxPlotR was used to generate box plots (Spitzer et al., 2014), in which the center lines represent the medians, box limits indicate 25th and 75th percentiles, and whiskers, determined by using the Tukey method, extended to data points <1.5 interquartile ranges from the first and third quartiles as determined by the BoxPlotR software.

## Acknowledgments

We thank Andrea Brand (Gordon Institute, University of Cambridge, England), Hiroshi Kimura (Tokyo Inst. of Technology, Tokyo, Japan), and Bloomington Stock Center for providing essential fly lines. We also thank Developmental Studies Hybridoma Bank (DSHB) for antibodies, and FlyBase for important genomic data. We are grateful for Eric Joyce (University of Pennsylvania, US) for providing us with the oligopaint probes and protocols. We thank the Nancy&Stephen Grand Israel National Center for Personalized Medicine (G-INCPM), and especially Eviatar Weizman, for processing and analyzing the DamId experiments. We thank R. Rotkopf from the Life Sciences Core Facility, Weizmann Institute, for statistical analysis. The images in this paper were acquired at the Advanced Optical Imaging Unit, de Picciotto-Lesser Cell Observatory unit, at the Moross Integrated Cancer Center Life Science Core Facilities, Weizmann Institute of Science. This study was supported by grants from “The French Muscular Dystrophy Association (AFM-Téléthon)” grant # 22339, and grant # 24142, and Israel Science Foundation (ISF) grant # 750/17. S. Safran is grateful for the support of the Volkswagen Foundation and the Perlman Family Foundation.

## Supplementary Information

**Figure S1:**
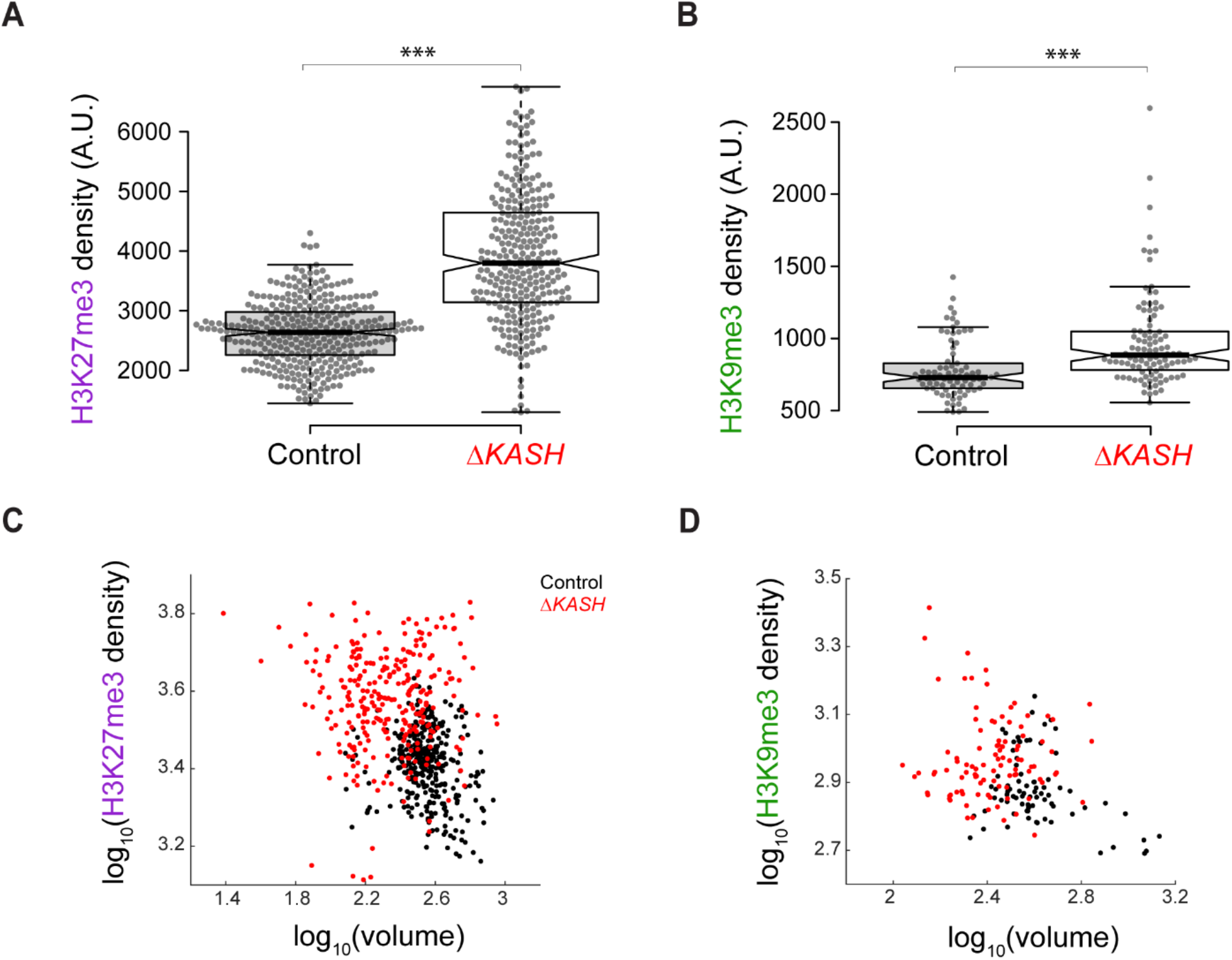
Increased repressive H3K27me3 and H3K9me3 chromatin density, inversely correlated with nuclear volume, in muscle fibers deficient of both *Msp300* and *klar* (ΔKASH). Quantification of mean nuclear fluorescence intensity shows increased **(A)** H3K27me3 (48% increase, p<0.001) and **(B)** H3K9me3 (25% increase p<0.001) in ΔKASH deficient muscle fibers. **(C, D)** Mean nuclear fluorescence intensity is plotted against the corresponding nuclear volume (log10 scale) for each epigenetic mark. **(A, C)** N=5 larvae, n=337 nuclei in control, N=5 larvae, n=284 nuclei in ΔKASH. **(B, D**) N=5 larvae, n=78 nuclei in control, N=4 larvae, n=100 nuclei in ΔKASH. For statistical significance ***p<0.001.

**Figure S2:**
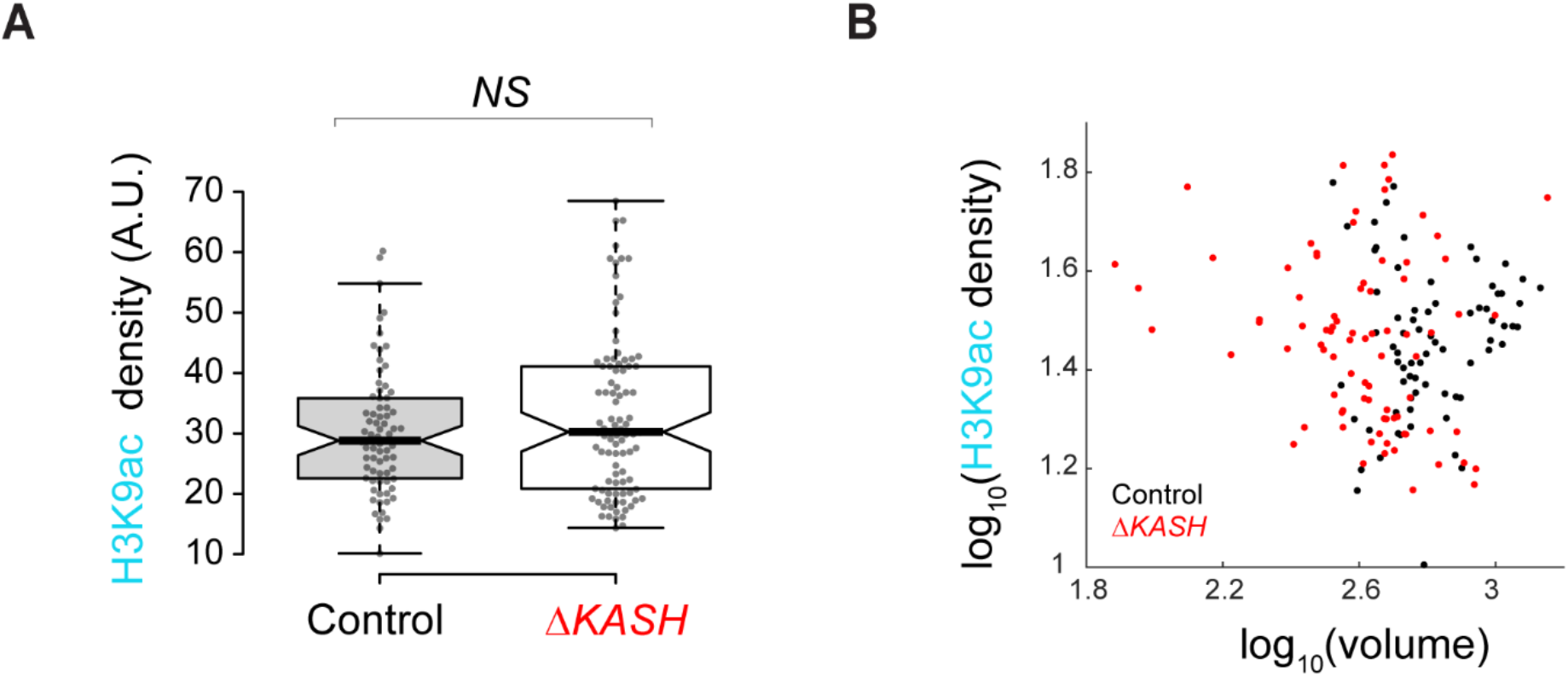
No change in active H3K9ac chromatin density in muscle fibers deficient of both *Msp300* and *klar* (ΔKASH). **(A)** Quantification of mean nuclear fluorescence intensity shows no change in H3K9ac in ΔKASH deficient muscle fibers (p=0.19). **(B)** No correlation in mean nuclear H3K9ac fluorescence intensity and the corresponding nuclear volume (log10 scale). N=5 larvae, n=73 nuclei in control, N=4 larvae, n=97 nuclei in ΔKASH.

**Figure S3:**
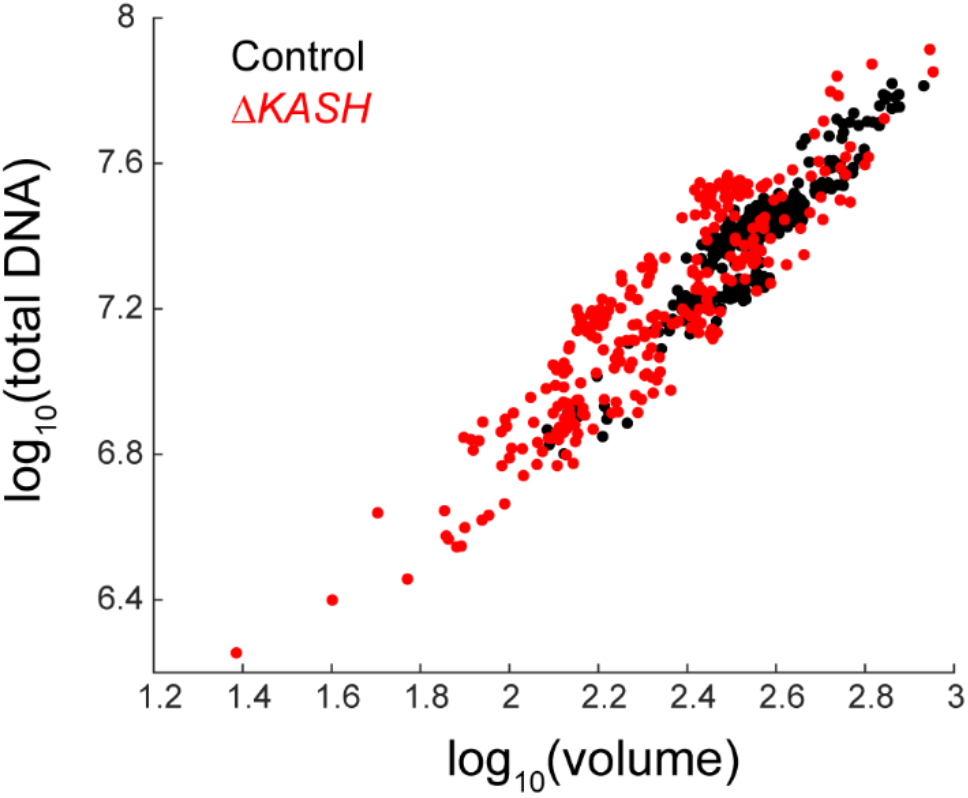
Preserved meso-scale DNA condensation in muscle fibers deficient of both *Msp300* and *klar* (ΔKASH). Total DNA intensity plotted versus nuclear volume (log10 scale) for ΔKASH (red) and control (black).

**Figure S4:**
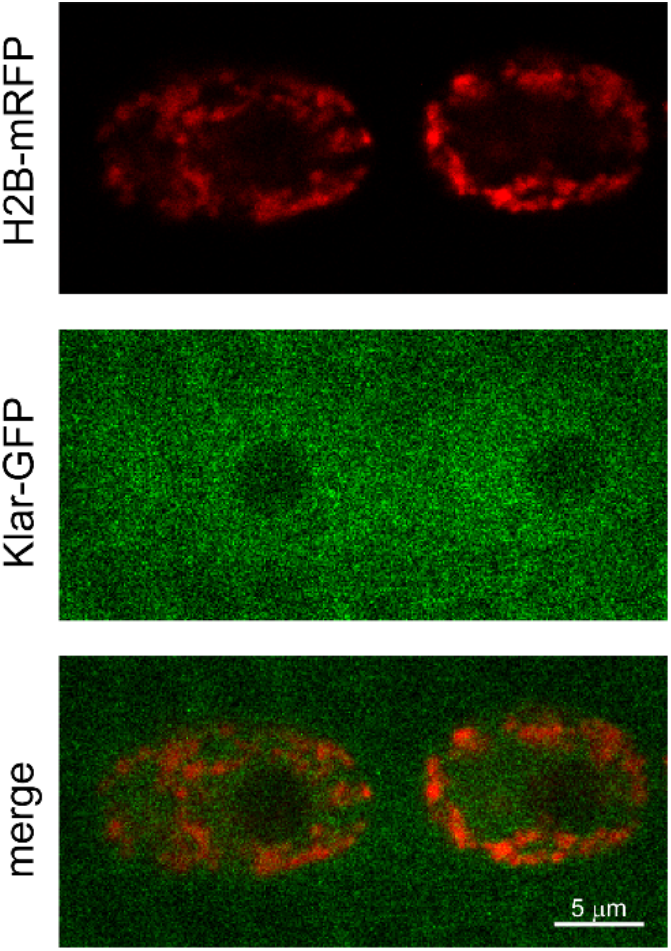
Live imaging of chromatin distribution in *SUN/koi* mutant muscle nuclei. Middle confocal section of muscle nuclei co-labeled with H2B-mRFP, and Klar-GFP. Peripheral chromatin distribution in the *SUN/koi* mutant can be observed from the H2B-mRFP channel. Klar-GFP fails to localize to the nuclear envelope in the *SUN/koi* mutant.

---

**<u>Movie S1:</u> 3D spatial distribution of H3K27me3-GFP in live, in-vivo control muscle nucleus**. 3D reconstruction from confocal z-stacks acquired with 0.308 μm intervals.

**<u>Movie S2:</u> 3D spatial distribution of H3K27me3-GFP in live, in-vivo***SUN/koi* **mutated muscle nuclei**. 3D reconstruction from confocal z-stacks acquired with 0.308 μm intervals.

